# Mapping the MIF-2 Chemokine Interactome Reveals MIF-2–CCL20 Complex Formation in Liver Fibrosis

**DOI:** 10.64898/2026.03.02.708954

**Authors:** Adrian Hoffmann, Markus Brandhofer, Yufei He, Genta Bushati, Elena Siminkovitch, Xavier Blanchet, Michael K. Otabil, Lin Zhang, Simon Ebert, Mathias Holzner, Christine Krammer, Kathleen Hille, Christian Wichmann, Hanno Niess, Omar El Bounkari, Christian Weber, Aphrodite Kapurniotu, Philipp von Hundelshausen, Patrick Scheiermann, Jürgen Bernhagen

**Affiliations:** Division of Vascular Biology, Institute for Stroke and Dementia Research (ISD), LMU University Hospital, Ludwig-Maximilians-Universität (LMU) Munich, Munich, Germany; Department of Anaesthesiology, LMU University Hospital, Ludwig-Maximilians-Universität (LMU) Munich, Munich, Germany; German Centre of Cardiovascular Research (DZHK), partner site Munich Heart Alliance, Munich, Germany; Institute for Cardiovascular Prevention, LMU University Hospital, Ludwig-Maximilians-Universität (LMU) Munich, Munich, Germany; Division of Peptide Biochemistry, TUM School of Life Sciences, Technische Universität München (TUM), Freising, Germany; Division of Transfusion Medicine, Cell Therapeutics and Haemostaseology, LMU University Hospital, Ludwig-Maximilians-Universität (LMU) Munich, Munich, Germany; German Cancer Consortium (DKTK), partner site Munich, Germany; Biobank of the Department of General, Visceral and Transplant Surgery, LMU University Hospital, Ludwig-Maximilians-Universität (LMU) Munich, Munich, Germany

**Keywords:** D-dopachrome tautomerase, D-DT, MIF-2, macrophage migration inhibitory factor, MIF, atypical chemokine, classical chemokine, CXCR4, liver fibrosis, chemokine heterocomplex, protein-protein interactions

## Abstract

D-dopachrome tautomerase (D-DT/MIF-2) is an inflammatory cytokine, atypical chemokine (ACK) and member of the macrophage migration-inhibitory factor (MIF) family. While interactions among classical chemokines (CKs) are established, ACK–CK interactions remain underexplored. Here, we screened for MIF-2 binding-partners using a protein array encompassing all CKs and selected ACKs, and validated candidate interactors by surface-plasmon resonance. CCL20/MIP-3α was prioritized based on RNA-sequencing suggesting induction during liver fibrosis. MIF-2/CCL20 complex formation was verified by microscale thermophoresis and interaction interfaces mapped using peptide array and *in-silico* modeling. The deduced binding-site near the MIF-2 tautomerase pocket was consistent with inhibition of its tautomerase activity by CCL20. We found both proteins abundantly expressed in human liver tissue, with a positive correlation. Pull-down confirmed complex formation and proximity ligation assay demonstrated MIF-2/CCL20 complexes in liver *in situ,* with higher levels in fibrotic tissue. Functionally, MIF-2/CCL20 complexes suppressed MIF-2–driven CD4⁺ T-cell chemotaxis and fibroblast IL-6 secretion, indicating modulation of immune and stromal responses. This study extends the ACK interactome to MIF-2 and suggests ACK/CK complexes modulate chemokine activities in liver fibrosis.

## Introduction

Failure to resolve inflammation can lead to organ fibrosis, a pathological process characterized by excessive extracellular matrix deposition and tissue dysfunction, representing a hallmark of end-stage disease that can affect any organ (Gilgenkrantz *et al*, 2025; Rockey *et al*, 2015; Zhao *et al*, 2022). While the molecular cues that govern the switch from resolution of inflammation to tissue fibrosis remain incompletely understood, chemokines represent promising therapeutic targets due to their upstream role as initiators, amplifiers, and sustainers of inflammatory signaling and leukocyte recruitment (Xu *et al*, 2023). However, chemokine-directed therapies remain underexploited, largely due to the system’s complexity and redundancy, with a multitude of ligands and receptors showing extensive cross-binding and context-specific functions. Classical chemokines (CKs) are categorized into several subclasses based on the spacing of conserved N-terminal cysteines, and can exhibit homeostatic (constitutive), inflammatory (inducible), or mixed-type functions (Xu *et al*., 2023). While CKs typically form homodimers to engage their cognate receptors, certain combinations such as CCL5/CXCL4 or CCL17/CCL5 can assemble into CC/CXC- or CC/CC-type heterocomplexes that alter the receptor binding and inflammatory or chemotactic properties, adding combinatorial complexity to the chemokine network on a systems level (von Hundelshausen *et al*, 2017). Atypical chemokines (ACKs), defined as protein mediators with chemokine-like properties but lacking CK-defining structural motifs, add an intriguing additional regulatory layer by modulating chemokine pathways through non-cognate engagement of chemokine receptors (Kapurniotu *et al*, 2019). Macrophage migration-inhibitory factor (MIF) is a prototypical ACK that signals via CXCR2, CXCR4, and ACKR3, in addition to its cognate receptor CD74, thereby influencing inflammatory responses in acute and chronic inflammation, fibrosis, and cancer (Bernhagen *et al*, 2007; Calandra & Roger, 2003; Kapurniotu *et al*., 2019; Leng *et al*, 2003; Sun *et al*, 2026; Weber *et al*, 2008). Similar to other pleiotropic inflammatory mediators such as IL-6, MIF has been reported to exert both pro- and antifibrotic effects depending on the tissue and disease context (Djudjaj *et al*, 2017; Heinrichs *et al*, 2021; Heinrichs *et al*, 2011; Luo *et al*, 2021; Miller *et al*, 2008; Wang *et al*, 2024a). In liver fibrosis, MIF mediates anti-fibrotic effects in toxin-induced injury, likely through CD74/AMP-activated protein kinase (AMPK) signaling in resident cells, whereas its pro-fibrotic activity in non-alcoholic steatohepatitis (NASH) appears to involve immune modulation, including the polarization of NKT cells toward a fibrosis-promoting phenotype (Heinrichs *et al*., 2021; Heinrichs *et al*., 2011).

D-dopachrome tautomerase (D-DT/MIF-2) is a structural homolog of MIF that displays distinct regulatory functions and a characteristic tissue-specific expression pattern, positioning it as an independent modulator of inflammatory processes in various disease settings (El Bounkari *et al*, 2025; Merk *et al*, 2011; Tilstam *et al*, 2021; Zan *et al*, 2022).

MIF-2 lacks the pseudo-ELR motif present in MIF and therefore does not bind CXCR2. However, MIF-2 has been implicated in engaging ACKR3 in chronic lung disease, and we previously identified MIF-2 as an ACK that induces CXCR4-dependent leukocyte chemotaxis with even greater potency than MIF (El Bounkari *et al*., 2025; Song *et al*, 2021). In the same study, MIF-2 was established as an independent risk factor for atherosclerosis, with elevated levels in unstable human plaques, correlation with coronary artery disease severity, and promotion of hepatic lipid accumulation in a murine high-fat diet model revealing a unique pro-lipogenic activity not observed for MIF (El Bounkari *et al*., 2025). The liver also displays the highest basal expression of MIF-2, pointing to an organ-specific role.

Building on the concept of the chemokine interactome and our prior identification of functional MIF–CXCL4L1 binding as a prototypical ACK/CK complex (Brandhofer *et al*, 2022), we asked whether MIF-2 also engages in heterocomplex formation with classical chemokines. We systematically mapped the MIF-2/chemokine interactome using a comprehensive chemokine protein array, followed by structural and biochemical validation of candidate interaction partners. Among these, CCL20 emerged as a high-affinity binding partner of MIF-2 and was prioritized for further exploration, also because it had previously been identified as part of a 25-gene fibrosis-progression signature in non-alcoholic fatty liver disease (NAFLD) (Govaere *et al*, 2020). We re-analyzed the corresponding RNA-seq dataset (GSE135251), confirmed progressive upregulation of CCL20 in advanced fibrosis stages, alongside abundant hepatic expression of MIF-2, and studied the hepatic formation of MIF-2–CCL20 complexes and its functional consequences for liver fibrosis, to unravel context-dependent fibrogenic hepatic chemokine activities.

## Results

### Discovery of the MIF-2 chemokine interactome by chemokine protein array and surface plasmon resonance

The identification of the chemokine interactome, i.e. the formation of heteromeric complexes between CKs from the same or even different sub-classes has revealed an emerging structural and functional complexity in the chemokine network, likely serving to fine-tune chemokine activities. With the discovery of heteromeric MIF/CXCL4L1 complexes (Brandhofer *et al*., 2022), we showed that this intriguing principle can even extend to interactions between ACKs (lacking chemokine structural folds) and CKs. MIF-2 exhibits only 34% sequence identity to MIF, but features an overall very similar three-dimensional architecture. We therefore hypothesized that MIF-2 might be able to engage in heteromeric interactions with certain CKs as well. To begin to test this notion, we performed a comprehensive unbiased chemokine protein array screen. The array comprised 47 human chemokines from all four structural subfamilies (CC, CXC, CX_3_C, and XC), along with selected damage-associated molecular pattern (DAMP)-type and ACK mediators including high-mobility group binding-1 (HMGB1), human β-defensins (HBDs), and peroxiredoxins (**Fig. 1A–B**) (He *et al*, 2019; Mantonico *et al*, 2024). The array was probed with biotin-conjugated human MIF-2, and binding partners were visualized by streptavidin–peroxidase-based detection **(Fig. 1A)**. The experiment displayed in **Fig. 1B** was performed at pH 8.0, and yielded the most defined interaction signals for MIF-2. A corresponding array carried out at pH 6.0 showed overall weaker and less distinct interaction patterns and was therefore not used as the primary reference condition (**Supp. Fig. 1A**). Several well-defined signal spots indicated robust binding of MIF-2 to multiple CKs, most prominently the CC-type chemokines CCL20, CCL24, CCL25, CCL26, and CCL28, as well as the CXC-type chemokines CXCL9, CXCL11, and CXCL17 (**Fig. 1B**). Additional weaker signals were detected for CCL21, XCL1, CXCL12α, and CXCL13, while other arrayed CKs showed negligible or no reactivity. Interestingly, a distinct signal was also observed for peroxiredoxin 1 (Prx1), suggesting a potential non-chemokine interactor, while HMGB and HBD proteins were negative. Confirming the validity of the chemokine array approach, strong positive spots were obtained for immobilized biotinylated MIF and MIF-2. Collectively, these data suggested that MIF-2 selectively interacts with a defined subset of CKs.

**Figure 1:**
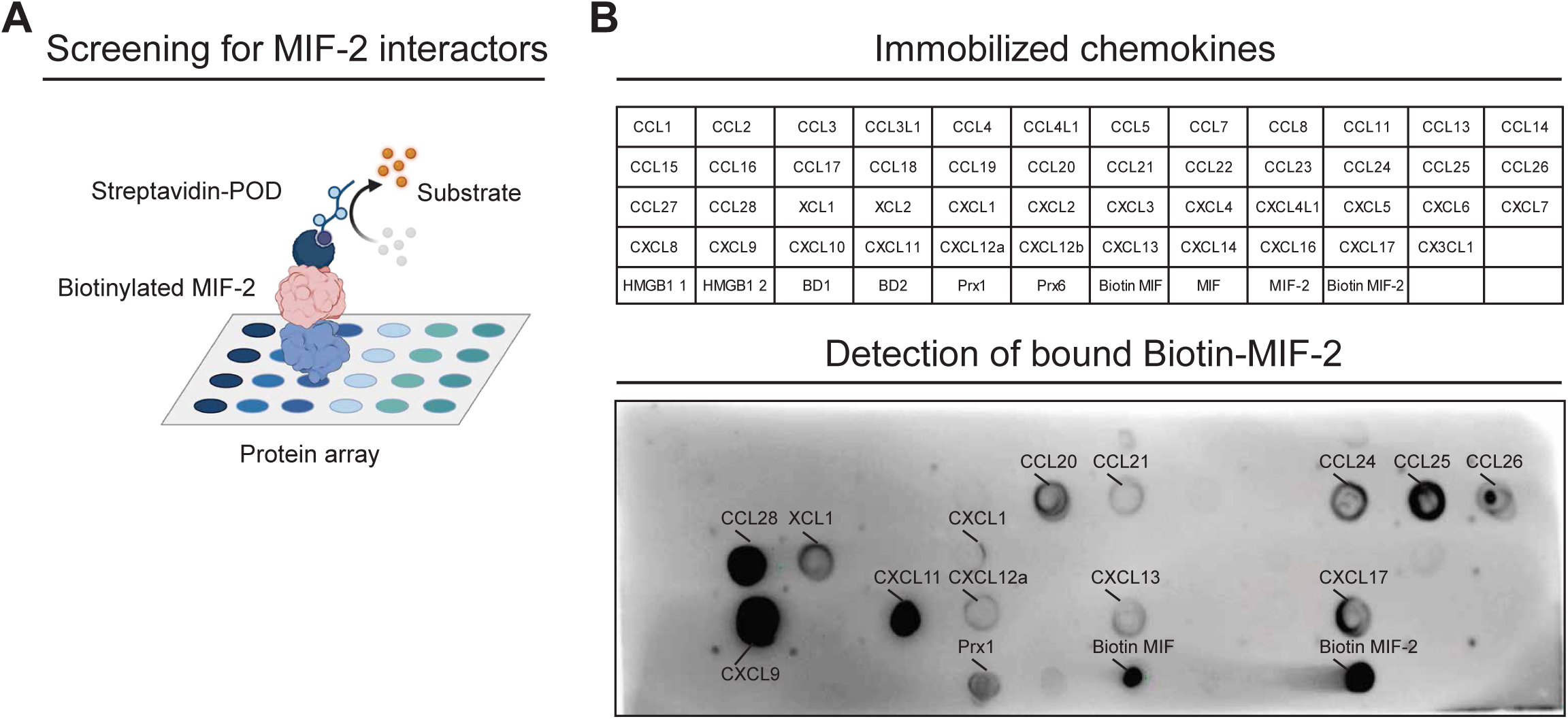
Unbiased protein-array analysis maps the MIF-2 interactome and uncovers selective binding to classical chemokines (CKs) and inflammatory mediators. (**A**) Schematic depiction of the experimental setup used to screen for chemokine-related MIF-2 interactors, in which CKs and atypical chemokines (ACKs) were immobilized on a nitrocellulose membrane, probed with biotin-MIF-2. Subsequently, bound biotin-MIF-2 was revealed by chemiluminescence. (**B**) Layout showing the position of CKs, ACKs, and selected inflammatory mediators on the nitrocellulose membrane (top), and the corresponding membrane developed against bound biotin-MIF-2 after incubation at pH 8.0 (bottom), revealing multiple candidate interactors.

To validate the candidate interactions identified in the chemokine array and determine the binding affinities, we next performed surface plasmon resonance (SPR) analysis using a MIF-2–coupled sensor chip. The most prominent candidate CKs were injected in the soluble phase at increasing concentrations, with CCL2 included as a negative control. Sensorgrams revealed concentration-dependent binding of multiple CKs to immobilized MIF-2, confirming their specific interaction (**Fig. 2A–B**). Quantitative evaluation showed that three CKs bound MIF-2 with high affinity K_D_ values in the sub-100 nM range: CCL20 (K_D_ 88.7±0.96 nM), CCL25 (K_D_ 70.9±25.3 nM), and CXCL17 (K_D_ 45±27.1 nM). Intermediate affinities between 100 and 1000 nM were observed for CXCL9 (K_D_ 106±13 nM), CXCL12 (K_D_ 258±61.1 nM), CXCL13 (K_D_ 205±78 nM), and CXCL11 (K_D_ 551±366 nM). The strongest binding was detected for CCL26 (K_D_ 2.0±1.28 nM) and CCL28 (K_D_ 1.15±1.81 nM). In contrast, CCL2 displayed only minimal binding (K_D_ 2460±1060 nM), indicative of no specific interaction in this assay. Taken together, the quantitative SPR approach independently confirmed the MIF-2/chemokine interactions suggested by the chemokine array and revealed low-nanomolar binding affinities for several pairs, consistent with the formation of stable heterotypic complexes between MIF-2 and selected CC and CXC chemokines.

**Figure 2:**
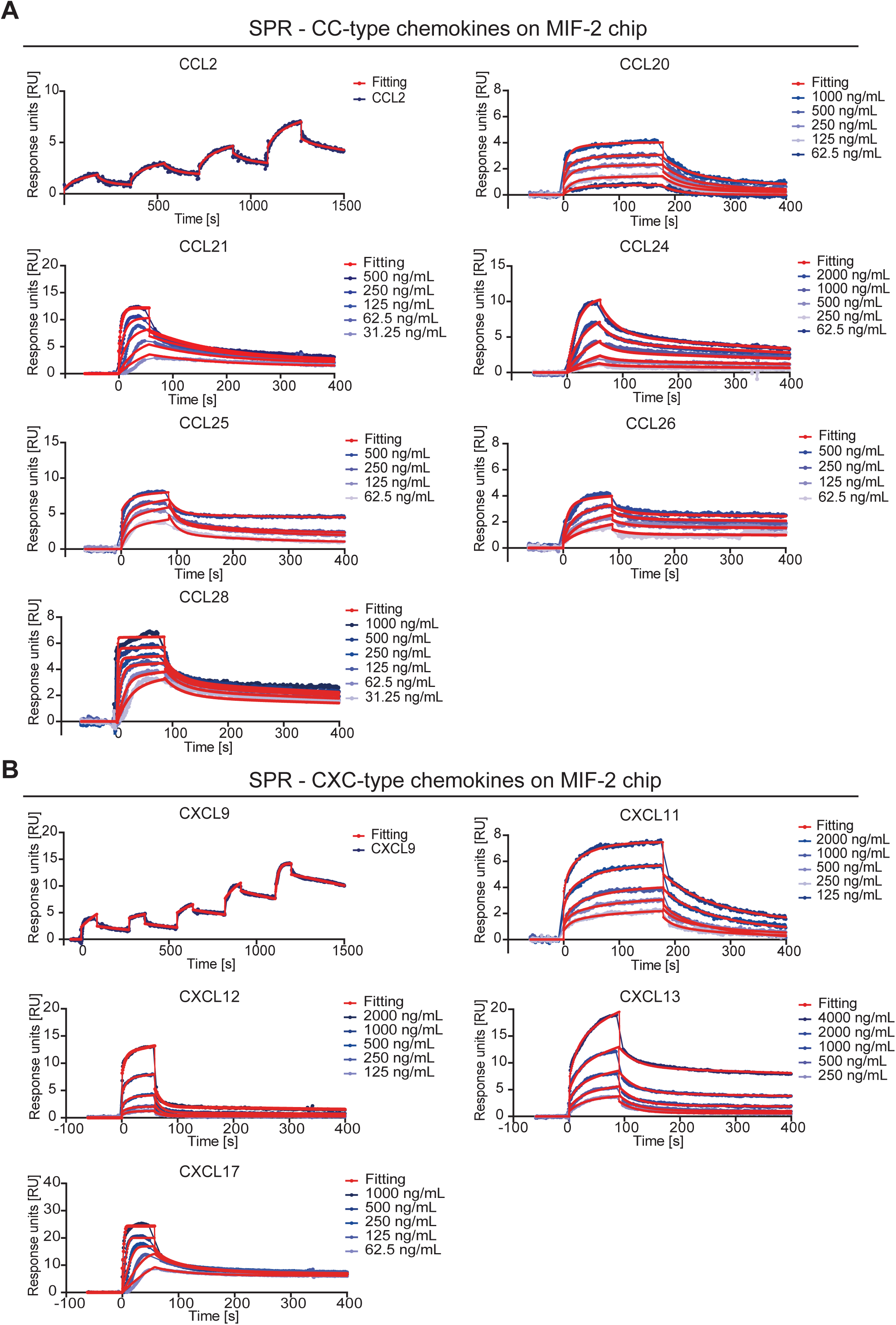
Surface plasmon resonance (SPR)-based validation of MIF-2 complex formation with selected candidate interactors. Depicted are sensograms of SPR experiments in which increasing concentrations of (**A**) CC-type chemokines and (**B**) CXC-type chemokines in the soluble phase were injected onto sensor-chips coated with biotin-MIF-2. K_D_ values for the tested interactions were in the low- to intermediate nanomolar range except for CCL2, which served as negative control based on the protein array screen. For CCL2, a K_D_ of 2.46 µM was calculated indicating markedly lower binding affinity compared with other MIF-2 interactors.

### Hepatic and fibrosis expression pattern prioritizes CCL20/MIF-2 interaction for further study

To prioritize for predictably biologically relevant interactions, we next considered the tissue localization and expression characteristics of MIF-2 and the confirmed suggested candidate interactors, in addition to a nanomolar binding affinity. Both MIF-2 and CCL20 are prominently expressed in the liver under healthy/homeostatic conditions and CCL20 has been amply implicated in hepatic inflammation and fibrosis (Chu *et al*, 2018; Govaere *et al*., 2020; Hanson *et al*, 2019). MIF-2 shows one of its highest basal expression levels in hepatic tissue, where it was previously identified by us to promote lipid accumulation in a high-fat-diet-based cardiovascular mouse model (El Bounkari *et al*., 2025). CCL20 was identified by Govaere et al as part of a 25-gene fibrosis-progression signature in human non-alcoholic fatty liver disease (NAFLD, now termed metabolic dysfunction–associated steatotic liver disease; MASLD), based on transcriptomic profiling of 206 liver biopsies spanning the full histological spectrum from NAFL to NASH with fibrosis stages F0–F4. (Govaere *et al*., 2020). To directly compare MIF-2 and CCL20 expression patterns in conjunction with their associated receptor network, we re-analyzed the available bulk RNA-sequencing data from the discovery cohort (GSE135251) of this study. Normalized expression values for MIF, MIF-2, CCL20 and their respective receptors — CD74 and CXCR4 as shared receptors for MIF and MIF-2, and CCR6 as the canonical receptor for CCL20 — were extracted and compared across disease stages (**Fig. 3**). This analysis confirmed the progressive up-regulation of CCL20 from NAFL to NASH and with increasing fibrosis severity (Govaere *et al*., 2020). By comparison, MIF and MIF-2 were abundantly expressed across all stages and showed only minimal changes with fibrotic progression. The only statistically significant difference was a modest down-regulation of MIF-2 between NAFL and NASH with fibrosis stage F2, while all other comparisons remained non-significant, consistent with a largely constitutive expression rather than disease-driven regulation. Analysis of receptor expression showed higher CXCR4 levels in fibrosis stages F2 and F3 compared with NAFL, whereas no significant difference was detected in F4, likely reflecting the smaller sample size and greater heterogeneity in this advanced stage. In contrast, CD74 showed consistently high expression levels without major changes across conditions. CCR6 was expressed merely at low levels, and no relevant changes were observed throughout the cohort (**Fig. 3**).

**Figure 3:**
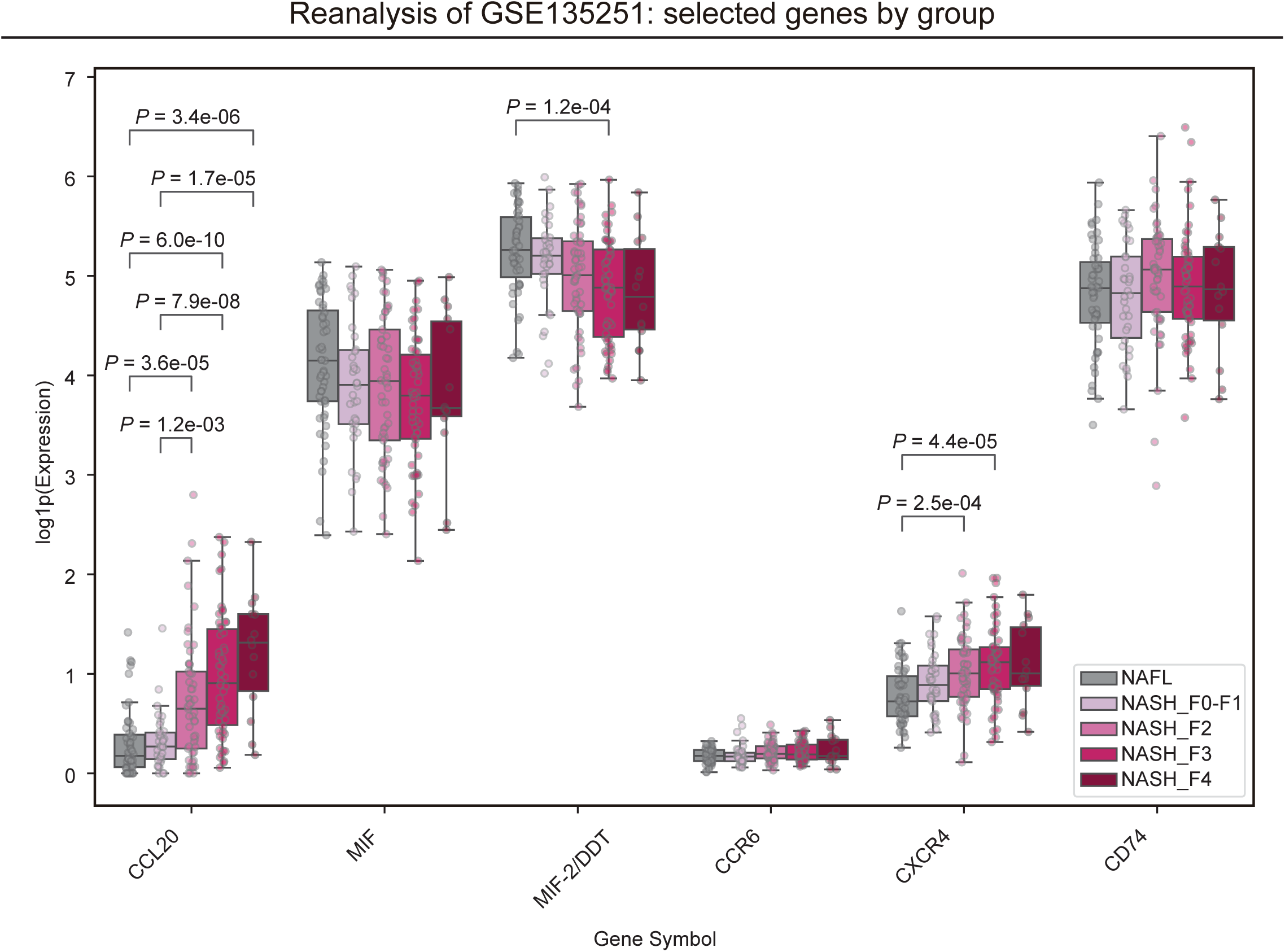
Hepatic expression patterns identify CCL20 as a prioritized MIF-2 interactor and implicate relevant receptor axes in liver fibrosis. Shown are gene expression data re-analyzed from the GSE135251 dataset, focusing on CCL20, MIF, and MIF-2 (DDT) together with their corresponding receptors CCR6, CXCR4, and CD74 in NASH across increasing fibrosis stages (F0–F4; shades of magenta) compared with NAFL (grey) (Govaere *et al*., 2020). Statistical comparisons between groups were performed using the Mann–Whitney U test, with *P* values adjusted for multiple testing using the Holm–Bonferroni correction. NASH: non-alcoholic steatohepatitis; NAFL: non-alcoholic fatty liver disease. Statistical significance is indicated by actual adjusted *P* values.

For comparison, we also assessed hepatic transcript levels of all other candidate interactors, including CCL21, CCL24, CCL25, CCL26, CCL28, CXCL9, CXCL11, CXCL12, CXCL13, CXCL17, XCL1 and PRDX1 (**Supp. Fig. 2**). Among these, CCL21 showed a stepwise increase with advancing fibrosis, similar to CCL20, whereas CCL28 was induced primarily between NAFL and NASH with fibrosis stage F4. In contrast, CCL24, CCL25, CCL26, CXCL9, CXCL11, CXCL12, CXCL13, CXCL17, XCL1 and PRDX1 did not show significant differential expression across fibrosis stages. In summary, our analysis revealed consistently high hepatic mRNA expression of MIF and MIF-2, with marked stage-dependent upregulation of CCL20. Receptor analysis showed uniformly low CCR6 expression, a fibrosis-associated increase in CXCR4 at intermediate stages, and stable high-level CD74 expression across conditions.

### Validation, structural mapping, and functional assessment of the MIF-2–CCL20 interaction by biophysical assays, *in silico* modeling, and tautomerase activity measurements

Based on the initial protein binding data and their expression characteristic in the liver and across different hepatic fibrosis stages, we prioritized the MIF-2/CCL20 pair for further scrutiny. To independently verify complex formation between MIF-2 and CCL20, we next applied microscale thermophoresis (MST), a technique that allows sensitive detection of molecular interactions in solution and has been previously validated in our laboratory for chemokine-binding analyses (Brandhofer *et al*., 2022; Kontos *et al*, 2020). In this setup, MST-red-labeled MIF-2 was titrated with CCL20. The titration yielded a dissociation constant of Kᴅ = 4958 ± 2000 nM (n=3) **(Fig. 4A)**. Although the Kᴅ was higher than that obtained by SPR, the characteristic sigmoidal curve characteristics clearly confirmed a specific interaction between MIF-2 and CCL20.

**Figure 4:**
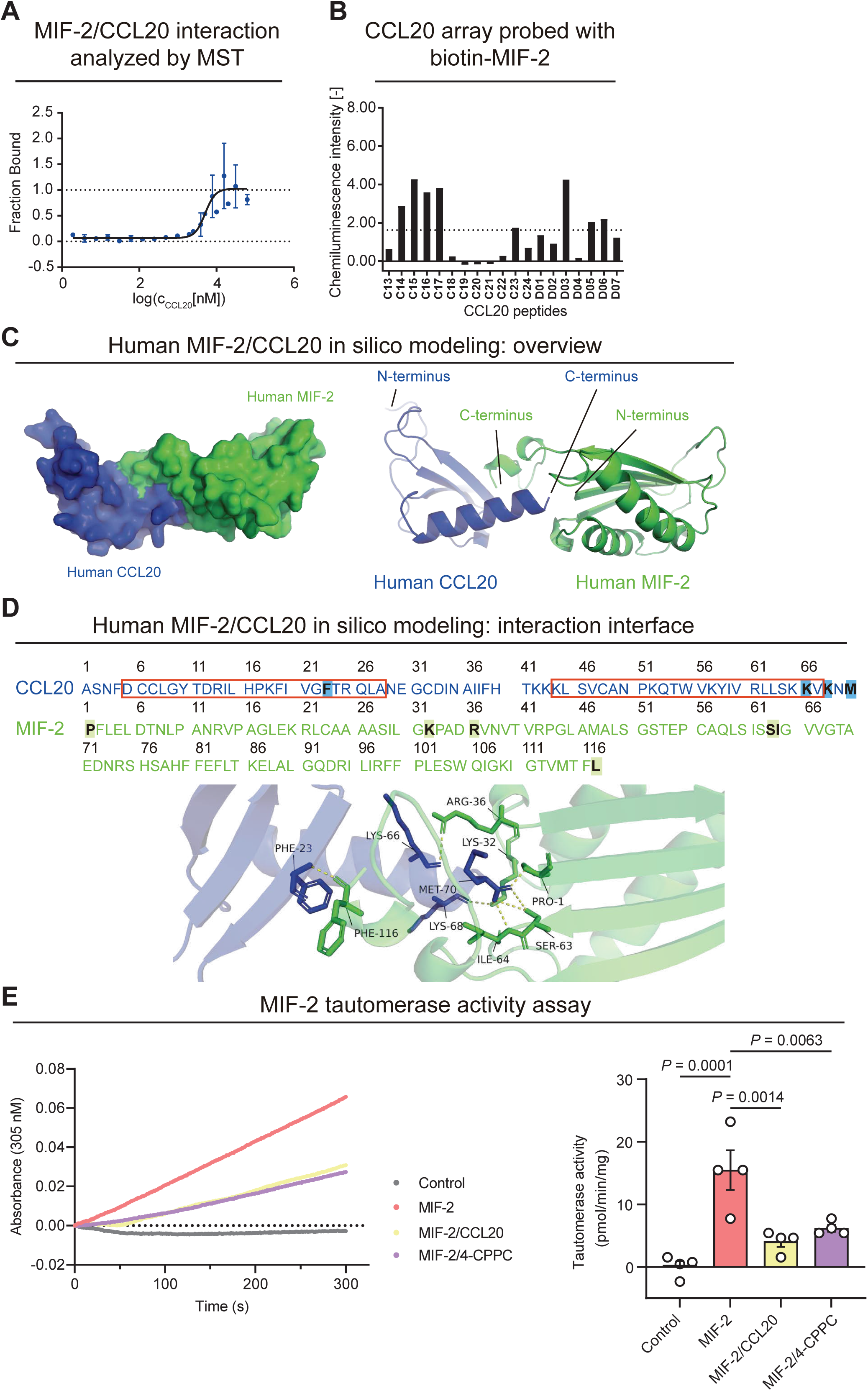
Characterization of MIF-2/CCL20 complex formation by microscale thermophoresis. (MST), peptide array analysis, *in silico* modeling and MIF-2 tautomerase activity assay. (**A**) Complex formation between fluorescently labeled MIF-2 and unlabeled CCL20 was validated in solution using MST, yielding a K_D_ of 5 ± 2 µM. Data are presented as mean ± standard deviation; n = 3. (**B**) CelluSpot peptide array analysis was performed to identify structural regions of CCL20 involved in MIF-2 binding. Overlapping CCL20-derived peptides were immobilized on the array and probed with biotinylated MIF-2. Chemiluminescence intensities reflect bound biotin–MIF-2. The dotted line marks the mean signal intensity across all CCL20 peptides. (**C**) Visualization of a heterodimer of human MIF-2 (blue) and CCL20 (green) predicted by AlphaFold 3.0. Shown on the left is a surface representation of the highest-scoring model; on the right, a cartoon rendering highlights the secondary structures. (**D**) Detailed depiction of the predicted interaction interface, alongside the AA-sequences of MIF-2 and CCL20. Residues predicted by AlphaFold 3.0 to contribute to protein–protein interactions are indicated in bold and framed in blue (MIF-2) or green (CCL20). Several of these residues lie adjacent to, or overlap with, the two CCL20 regions identified as potential MIF-2 binding sites in the CelluSpot peptide array shown in (**B**), highlighted here with red rectangles. (**E**) Representative kinetic curves of a fluorometric 4-hydroxyphenylpyruvate (HPP) tautomerization assay assessing MIF-2 tautomerase activity. Absorbance at 306 nm was monitored over time for reactions containing control buffer, 55 nM MIF-2 alone, or 55 nM MIF-2 in the presence of 30 nM 4-CPPC or 30 nM CCL20. Tautomerase activity was calculated from the initial linear phase of the reaction using an experimentally determined molar extinction coefficient (ε₃₀₆), with the corresponding quantification shown in the bar graph on the right. Data are shown as mean ± SD (n = 4). Statistical comparisons were performed using an unpaired Student’s t test. Statistical significance is indicated by actual *P* values.

In order to map the molecular interface of CCL20 involved in MIF-2 recognition, we generated a set of overlapping 15-mer peptides derived from the mature CCL20 sequence, each shifted by three amino acids (AA), thereby covering the entire CCL20 sequence (**Fig. 4B**). The peptides were immobilized on a glass array and probed with biotinylated MIF-2. This peptide-based screening for MIF-2 binding residues within CCL20 revealed a pronounced binding signal for a contiguous stretch of peptides (C14–C17), corresponding to the AA-sequence DCCLGYTDRILHPKFIVGFTRQLA, located immediately downstream of the conserved CC-motif in the N-terminal part of CCL20 (**Fig. 4B**). This segment coincides with the β-sheet region (“site 1”) of the canonical chemokine-fold described in the NMR structure of human CCL20 (PDB 1M8A). It is considered responsible for binding to the N-terminus of the receptor and thus is central to the two-site chemokine–receptor binding model and critical for CCR6 engagement (Crump *et al*, 1997; Hoover *et al*, 2002; Rajagopalan & Rajarathnam, 2006; Wasilko *et al*, 2020). Additionally, a pronounced signal was detected for peptide D03 (AA-sequence: KLSVCANPKQTWVKY) near the C-terminus of CCL20. Although the directly adjacent peptide D04 showed no detectable reactivity, weaker signals were observed for the partially overlapping peptides D05 and D06. Because D03, D05, and D06 together span the continuous C-terminal sequence KLSVCANPKQTWVKYIVRLLSKKV, we conclude this region is a second potential interaction site (**Fig. 4B**).

To model potential spatial relationships within a MIF-2/CCL20 complex, *in silico* heterodimer structures were generated using AlphaFold 3.0 (www.alphafoldserver.com) based on the mature forms of both proteins, with MIF-2 lacking the initiator methionine and CCL20 lacking the N-terminal signal peptide. AlphaFold generated five independent structural models (models 0–4), which all displayed a broadly similar overall complex architecture. Notably, across all five models, the C-terminal region of CCL20 consistently occupied the MIF-2 tautomerase pocket — an evolutionarily conserved site involved in MIF-2 receptor binding and a target for the discovery of small molecule MIF-2 inhibitors — with model-specific variations only noted in a few contributing residues. In line with this, the confidence estimate for the overall architecture—expressed as the predicted template modeling (pTM) score—indicated moderate global reliability (pTM = 0.65), while the metric that reflects confidence in the predicted interface geometry (the interface predicted TM-score (ipTM)) was low (ipTM = 0.32). The low ipTM denotes uncertainty in the predicted relative positions and interface orientation of the subunits and therefore warrants cautious interpretation. However, the recurring placement of the CCL20 C-terminus within the conserved MIF-2 pocket across all five models suggests that the overall interaction motif may be more stable than implied by the ipTM alone. Model 0 was selected for further visualization because it received the highest internal ranking among the generated predictions (**Fig. 4C–D**). This model was displayed both as a surface representation to illustrate the overall spatial arrangement of the complex and as a secondary-structure depiction to highlight fold-organization and the positions of the N- and C-termini (**Fig. 4C**). Closer inspection of model 0 enabled the predicted interface residues to be mapped onto the primary AA-sequences of MIF-2 and CCL20 (**Fig. 4D**). Individual residues within the predicted CCL20 interface corresponded to experimentally identified N-and C-terminal interaction regions obtained by the peptide array data, indicating qualitative convergence between computational and experimental mapping. Models 1–4 are provided in the **Supp. Figure 3** for comparison. Taken together, the AlphaFold predictions and peptide-array mapping jointly support a stable key–lock–type anchoring of the CCL20 C-terminus within the MIF-2 tautomerase pocket, despite computational variability at the residue and subunit-orientation levels. To functionally validate this structural prediction, we next assessed the impact of CCL20 on MIF-2’s vestigial tautomerase activity. In a fluorometric tautomerase assay, CCL20 significantly inhibited the catalytic activity of MIF-2. Of note, the inhibition profile and inhibitory potential was comparable to that of the established small-molecule MIF-2 inhibitor 4-CPPC, which directly occupies the same catalytic pocket (Pantouris *et al*, 2018; Tilstam *et al*, 2019) (**Fig. 4E, Supp. Fig. 4**). At equimolar concentrations, CCL20 reduced MIF-2 activity to levels nearly identical to those achieved with 4-CPPC, supporting the conclusion that both molecules act on the tautomerase functional interface.

Our combined biophysical analyses provide independent evidence for MIF-2/CCL20 complex formation. SPR and MST confirmed direct binding between MIF-2 and CCL20, while peptide-array profiling identified two CCL20 regions that may contribute to this interaction, consistent with *in silico* models positioning CCL20 within the MIF-2 tautomerase pocket. In line with this, CCL20 suppressed MIF-2 tautomerase activity with an efficacy comparable to the small molecule compound 4-CPPC. Together, these complementary approaches establish MIF-2/CCL20 complex formation and provide a structural framework for this non-canonical ACK–CK pairing.

### Biochemical and *in-situ* detection of MIF-2/CCL20 complexes in human liver and fibrosis

Building on the observed transcriptional expression patterns in NAFLD–NASH/fibrosis cohorts (see **Fig. 3**), we asked whether hepatic mRNA expression of the *MIF-2* and *CCL20* genes is reflected at the protein level, and whether MIF-2 and CCL20 form complexes *in situ*. To address this, we obtained clinically characterized human liver samples from the LMU-HTCR biobank and applied a complementary set of protein-based approaches, including quantitative immunoblotting, pulldown assays, and *in situ* PLA, to analyze MIF-2 and CCL20 protein co-expression and visualize MIF-2/CCL20 complexes in fibrotic and non-fibrotic liver tissue. Fibrosis status and corresponding patient characteristics were provided by the LMU-HTCR biobank, with fibrosis grading performed by a board-certified pathologist based on matched histological sections (**Supp. Table 1**).

First, we assessed MIF-2 and CCL20 protein expression in human liver tissue by SDS–PAGE/Western Blot (WB). Both proteins were readily detectable in all available samples and exhibited expected inter-individual variability (**Fig. 5A-B**). MIF-2 generally showed higher expression levels, consistent with the transcriptomic data confirming strong hepatic expression. Across all samples, MIF-2 and CCL20 levels displayed a significant positive correlation (R = 0.7727, *P* = 0.0074) suggesting coordinated regulation within the hepatic microenvironment (**Fig. 5C**). However, when comparing non-fibrotic (control) and fibrotic liver tissue, protein expression of both MIF-2 and CCL20 showed no significant differences (control n = 7, fibrosis n = 4; **Supp. Fig. 5A**). Despite the limited cohort size, the consistent detection of both proteins and their correlated expression patterns across samples support a stable hepatic presence of MIF-2 and CCL20.

**Figure 5:**
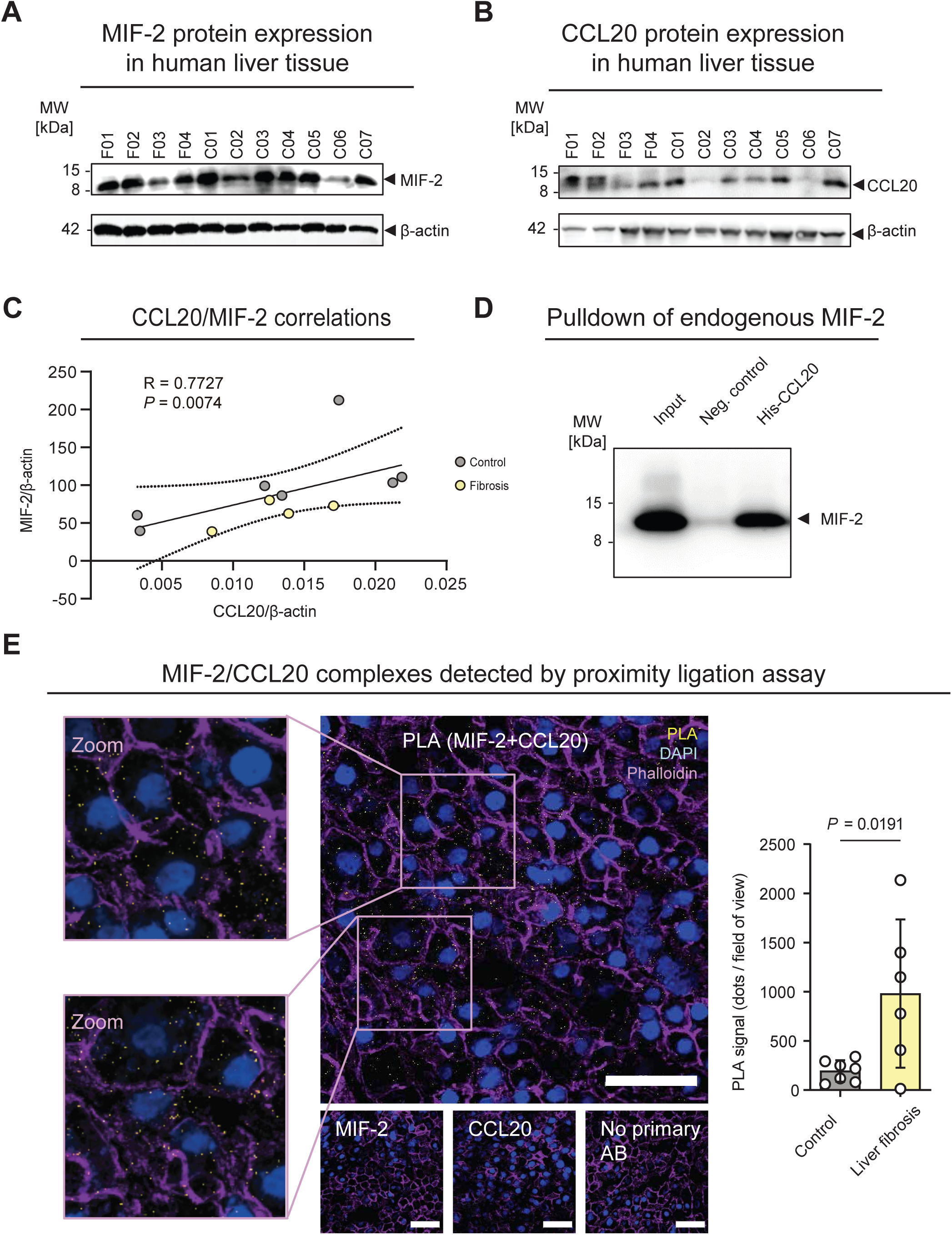
Abundant hepatic expression of MIF-2 and CCL20 and in situ detection of MIF-2/CCL20 heterocomplexes in human liver tissue. **(A)** Protein expression of MIF-2 in non-fibrotic control liver biopsies (coded “C…”) and fibrotic samples (coded “F…”) detected by SDS–PAGE/Western blot (WB) with β-actin serving as loading control. **(B)** Protein expression of CCL20 in the same samples shown in (**A**). **(C)** Spearman correlation analysis of MIF-2 and CCL20 protein levels normalized to the corresponding β-actin expression from (**A**) and (**B**). **(D)** Semi-endogenous pull-down assay in which recombinant His-tagged CCL20 was used to capture endogenous MIF-2 from human liver tissue lysates. Complexes were precipitated using NTA–cobalt paramagnetic beads with high affinity for polyhistidine tags. A representative blot of three independent experiments is shown, probed for MIF-2. “Input” refers to untreated tissue lysate prior to pull-down. “Neg. control” denotes a pull-down performed without addition of His-CCL20 (beads + lysate only), whereas “His-CCL20” indicates successful pull-down of endogenous MIF-2 upon addition of His-CCL20. **(E)** *In situ* visualization of MIF-2/CCL20 heterocomplexes in fibrotic human liver tissue by Proximity Ligation Assay (PLA). PLA-positive signals (yellow) indicate close proximity between MIF-2 and CCL20, consistent with heterocomplex formation. Tissue was counterstained with phalloidin (purple), and nuclei were stained with DAPI (blue). Negative controls (MIF-2 antibody only, CCL20 antibody only, or omission of primary antibodies) are shown in the inset panels. Fibrotic (n = 6) and non-fibrotic (n = 7) samples were analyzed with ≥4 fields of view per sample; quantification of PLA signal counts is shown in the graph on the right. Images were acquired by multiphoton microscopy; scale bar: 50 µm. Statistical comparisons were performed using an unpaired Student’s t-test, and data are presented as mean ± SD. Statistical significance is indicated by actual *P* values.

Given the higher hepatic protein expression of MIF-2, we next performed semi-endogenous co-immunoprecipitation using recombinant His-tagged CCL20 to pull-down endogenous MIF-2 from human liver lysates. Immunoblotting for MIF-2 revealed a clear band in CCL20 pull-downs, whereas no signal was detected in control pull-downs, in which lysates were incubated with paramagnetic beads alone (i.e., without His-CCL20), confirming assay specificity (**Fig 5D**, n = 5). To further substantiate these findings, we performed a complementary “vice-versa” pull-down using recombinant proteins, in which biotinylated MIF-2 immobilized on streptavidin-coated paramagnetic beads successfully co-precipitated soluble CCL20, thereby validating the bidirectional capacity of MIF-2 and CCL20 to form a complex (**Supp. Fig. 1B**). Finally, we employed PLA, a highly sensitive and widely used method for detecting protein–protein interactions *in situ*. PLA enables visualization of endogenous protein proximity by generating fluorescent signals only when target proteins lie within < 40 nm (Alam, 2022). PLA was performed on human liver tissue obtained from the HTCR biobank (non-fibrotic/control n = 7; fibrotic n = 6). After establishing specificity of antibody co-staining conditions (**Supp. Fig. 5B**), PLA was applied to assess MIF-2/CCL20 complex formation within the tissue architecture, and PLA signals quantified for each patient sample. Distinct PLA signals indicative of close molecular proximity between MIF-2 and CCL20 were readily detected across specimens. Notably, PLA signal frequency was increased in fibrotic compared with non-fibrotic liver tissue, suggesting an enhanced likelihood of MIF-2/CCL20 complex formation in liver fibrosis (**Fig. 5E; Supp. Table 1**).

Collectively, these data demonstrated that MIF-2 and CCL20 are co-expressed and form complexes that are enriched in fibrotic human liver tissue, underscoring their potential relevance in hepatic inflammation and fibrosis-associated tissue remodeling.

### Functional impact of MIF-2/CCL20 complex formation on CD4⁺ T cell migration and fibroblast IL-6 secretion

To explore functional consequences of MIF-2/CCL20 complex formation, we next focused on CD4⁺ T cells, central coordinators of immune and fibrotic processes via cytokine release and stromal interactions (Al Sayegh *et al*, 2025; Gomes *et al*, 2016; Meng *et al*, 2012). To determine whether MIF-2 and its interaction with CCL20 could influence the recruitment of these cells into fibrotic environments, we assessed their migratory response using an established 3D chemotaxis assay that enables single-cell live-tracking within a tissue-mimetic collagen matrix, thereby resolving individual migration trajectories. This assay, previously validated for MIF-induced CD4⁺ T-cell chemotaxis (Zhang *et al*, 2024), provides a suitable system to dissect CXCR4-dependent effects, as non-activated CD4⁺ T cells highly express CXCR4 as their sole MIF receptor. MIF-2 was recently identified as an additional CXCR4-ligand, yet its migratory effects on T cells have not been characterized (El Bounkari *et al*., 2025). CCR6 is expressed by CD4⁺ T cell subsets such as Treg and Th17 cells, and the CCL20–CCR6 axis governs T-cell trafficking in disease (Affò *et al*, 2015; Ito *et al*, 2019; Liu *et al*, 2025; Wang *et al*, 2009; Wang *et al*, 2024b).

Cell migration was recorded over a 2-hour period with 1-minute intervals. Stimulation with MIF-2 (400 ng/ml) induced a significant chemotactic response, whereas CCL20 (300 ng/ml) elicited only a weak, non-significant directional migration, in line with the low proportion of CCR6⁺ CD4⁺ T cells (**Fig. 6A-B**, **Supp. Fig. 6A**) (Francis *et al*, 2008). Strikingly, simultaneous stimulation with MIF-2 and CCL20 significantly inhibited chemotaxis, indicating interference between MIF-2- and CCL20-driven migratory cues (**Fig. 6A-B)**. In direct comparisons, CD4⁺ T cells consistently migrated toward the single-chemokine stimulus and away from the MIF-2/CCL20 complex. When the complex was present on both sides of the gradient, no directed migration was observed, suggesting neutralization of chemotactic signaling.

**Figure 6:**
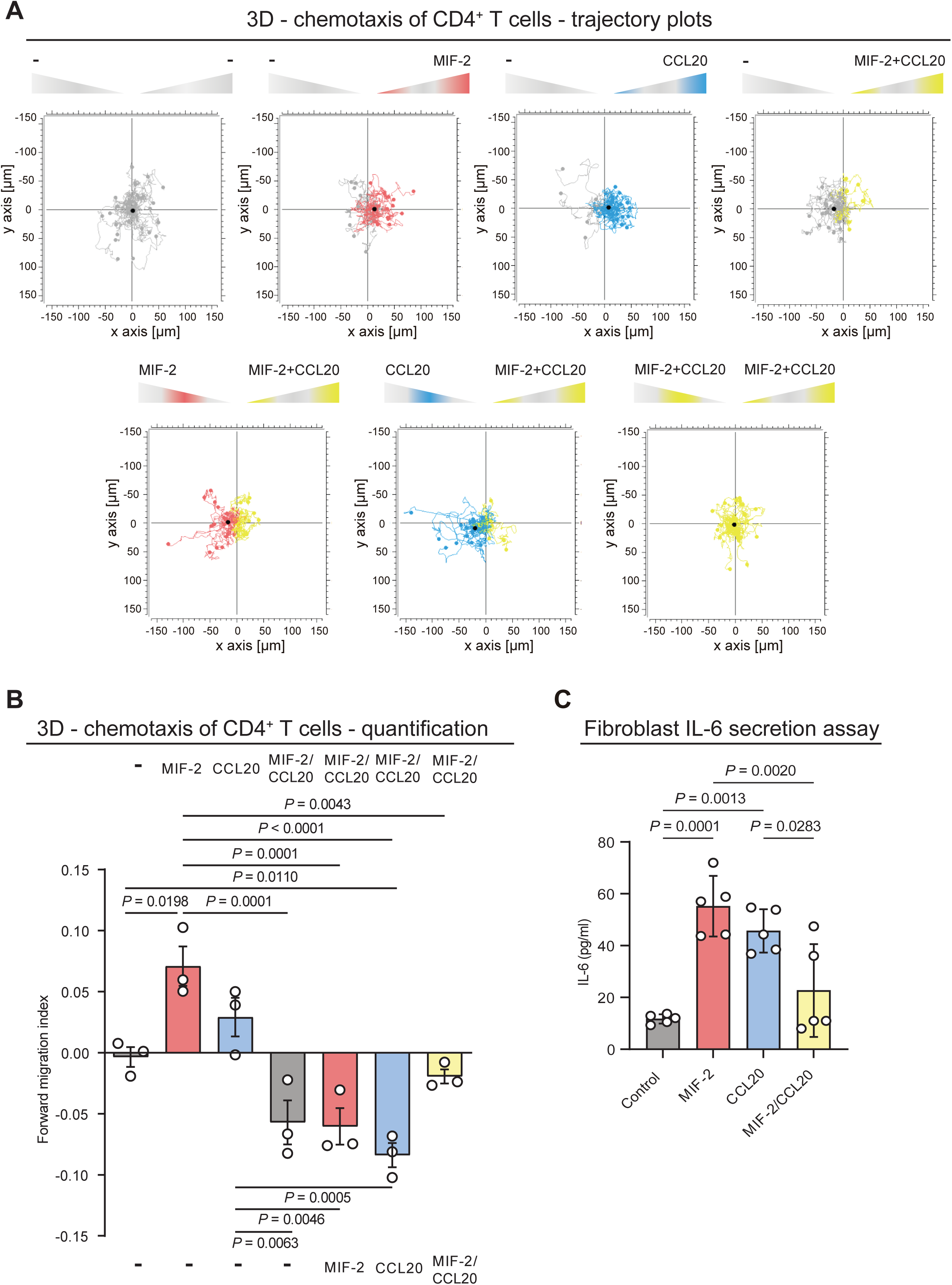
Functional inhibitory capacity of the MIF-2/CCL20 heterocomplex in human CD4⁺ T-cell migration and fibroblast IL-6 secretion. (**A**) Representative trajectory plots of migrating primary human CD4⁺ T cells exposed to gradients of MIF-2 (400 ng/ml), CCL20 (300 ng/ml), both combined, or control medium in an Ibidi µ-Slide 3D Chemotaxis assay. Treatments were added to the chemoattractant reservoirs on either side of the migration channel containing gel-embedded cells. Cell movement was recorded by live-cell time-lapse microscopy for 90 min at 37 °C, with 1-min intervals using a Leica DMi8 microscope, and individual migration tracks were reconstructed from the resulting image sequences. Tracks of cells migrating toward a given stimulus are color-coded (MIF-2: red; CCL20: blue; both: yellow), whereas control medium (grey) is indicated by “–”. The starting position of each cell was centered at x = y = 0; the black dot represents the center of mass of the cell population at the end of the experiment. (**B**) Quantification of the experiments shown in (**A**). The forward migration index (FMI) along the chemokine gradient (x-axis) is plotted, calculated from individual tracks of ≥30 cells per condition. Annotations above and below the diagram indicate the treatments applied to the right and left reservoirs, respectively, corresponding to the positive and negative x-axis in (A). n = 3. (**C**) IL-6 secretion by primary human fibroblasts stimulated with MIF-2 (1000 ng/ml), CCL20 (500 ng/ml), or both, measured by ELISA. Data are presented as mean ± SD. n = 5. Statistical analysis was performed by one-way ANOVA with Tukey’s post-hoc correction for **B** and **C**. Statistical significance is indicated by actual *P* values.

To further characterize the functional consequences of MIF-2/CCL20 complex formation beyond immune-cell migration, we next tested fibroblast activity. We stimulated primary human dermal fibroblasts with recombinant MIF-2 and CCL20 and quantified IL-6 secretion as an inflammatory readout via ELISA (**Fig. 6C)**. IL-6 is a central cytokine involved in fibroblast activation and myofibroblast differentiation and therefore served as an informative indicator of fibroblast inflammatory activity (Li *et al*, 2022).

Fibroblasts were selected as key effector cells in fibrotic disease, driving extracellular matrix deposition, tissue contraction, and chronic inflammation (Rockey *et al*., 2015). MIF is an established inducer of IL-6 in multiple cell types, including monocytes/macrophages and fibroblasts (Bernhagen *et al*, 1993; Santos *et al*, 2004; Zhang *et al*, 2021). MIF-2 has been reported to induce IL-6 mRNA expression in preadipocytes and to promote fibroblast viability and proliferation, while CCL20 induces IL-6 release in fibroblast-like synoviocytes (Alaaeddine *et al*, 2011; Ishimoto *et al*, 2012; Kim *et al*, 2017). However, whether MIF-2 elicits IL-6 release in fibroblasts had not been addressed prior to this study. The fibroblasts used here expressed low levels of CXCR4 (∼15%), no detectable CD74, and contained a substantial proportion of CCR6⁺ cells (∼50%) (**Supp. Fig.6 B-D**), providing a suitable system to assess potential effects on both the MIF-2/CXCR4 and CCL20/CCR6 axes. Stimulation with recombinant MIF-2 (1000 ng/ml) or CCL20 (500 ng/ml) each led to a significant increase in IL-6 secretion compared with unstimulated controls (control: 11.7 ± 1.77 pg/ml; MIF-2: 55.2 ± 11.7 pg/ml; CCL20: 45.6 ± 8.4 pg/ml). Strikingly, co-stimulation with MIF-2 and CCL20 markedly reduced IL-6 release relative to either ligand alone (MIF-2/CCL20: 22.7 ± 17.9 pg/ml), indicating that the combined presence of both chemokines counteracts their individual IL-6–inducing effects (**Fig. 6C**). These results reveal a functional crosstalk between MIF-2 and CCL20 in fibroblasts that dampens inflammatory cytokine output.

In summary, co-stimulation with MIF-2 and CCL20 inhibited directed CD4⁺ T-cell migration and suppressed fibroblast IL-6 secretion relative to either ligand alone, demonstrating an inhibitory effect of the MIF-2/CCL20 complex on the function of both individual mediators in immune and stromal cells.

## Discussion

Here, we define the MIF-2-interactome and identify CCL20 as a previously unrecognized high-affinity binding partner forming *in situ* complexes in human fibrotic liver tissue. Functionally, MIF-2/CCL20 complexes attenuate MIF-2-driven CD4⁺ T-cell migration and fibroblast IL-6 production, revealing an intrinsic modulatory role of this ACK–CK interaction in inflammation and fibrosis.

Fibrosis represents the maladaptive endpoint of a failed resolution process, in which chronic inflammation transitions into persistent tissue remodeling and dysfunction. This transition engages a coordinated multicellular network of immune and stromal cells and initiates inflammatory cascades that recruit leukocytes and promote fibroblast activation (Gilgenkrantz *et al*., 2025; Rockey *et al*., 2015; Zhao *et al*., 2022). The secretion of cytokines such as IL-1β, IL-6, TNF-α, TGF-β1, and IL-17A, together with multiple chemokines and growth factors, generates localized signaling domains that sustain pro-inflammatory and profibrotic responses (Chu *et al*., 2018; Gallucci *et al*, 2006; Govaere *et al*., 2020; Greenman *et al*, 2023; Hanson *et al*., 2019; Li *et al*, 2015; Meng *et al*., 2012; Sato-Matsubara *et al*, 2017; Xi *et al*, 2021).

The mechanisms determining whether inflammation resolves or evolves into fibrosis remain incompletely defined, reflecting complex context- and stage-dependent shifts in the inflammatory microenvironment, in which identical mediators can elicit divergent, even opposing, effects. These observations imply regulatory crosstalk mechanisms beyond a simple balance of pro- and anti-inflammatory mediators.

An under-studied mechanism is the fine-tuning of responses through protein–protein interactions among mediators. Chemokines assemble into homo- and heteromeric complexes via conserved interfaces to modulate receptor activation and downstream signaling, yet most interactions remain poorly defined (von Hundelshausen *et al*., 2017). CK interactions with ACKs have only recently emerged, with four examples reported to date, including CXCL12 complexes with HMGB1 or galectin-3 that differentially regulate CXCR4 signaling and HNP1 binding to CCL5 to promote CCR5-mediated monocyte adhesion (Alard *et al*, 2015; Eckardt *et al*, 2020; Kapurniotu *et al*., 2019; Mantonico *et al*., 2024; Schiraldi *et al*, 2012). Finally, we recently showed that the ACK MIF, acting as a non-canonical CXCR4 ligand, forms a complex with CXCL4L1, thereby suppressing MIF/CXCR4-driven leukocyte recruitment and thrombus formation (Brandhofer *et al*., 2022). Beyond CXCR4, MIF also signals via CD74, CXCR2, and ACKR3, exerting context-dependent pro- or antifibrotic effects (Djudjaj *et al*., 2017; Heinrichs *et al*., 2021; Heinrichs *et al*., 2011; Luo *et al*., 2021; Miller *et al*., 2008; Wang *et al*., 2024a). The MIF paralog MIF-2 is increasingly recognized as an independent and pathophysiologically relevant regulator and exhibits a particularly high basal expression in the liver (El Bounkari *et al*., 2025; Merk *et al*., 2011; Tilstam *et al*., 2021; Zan *et al*., 2022). Although MIF and MIF-2 share a similar but not identical three-dimensional fold they exhibit only ∼34% sequence identity and differ in key structural elements, including the absence of the pseudo-ELR motif in MIF-2, precluding CXCR2 engagement (Sugimoto *et al*, 1999; Tilstam *et al*., 2021). We recently characterized MIF-2 as an ACK that signals via CD74 and CXCR4 to promote pro-atherogenic leukocyte recruitment and metabolic inflammation, with *Mif-2* deficiency attenuating hepatic lipid accumulation (El Bounkari *et al*., 2025). Evidence for a contribution of MIF-2 in fibrosis remains scarce. In experimental heart failure, *Mif-2* deficiency aggravated interstitial fibrosis, potentially via TGF-β/Smad signaling, whereas upregulated hepatic MIF-2 levels in CCl₄-induced liver injury remained without mechanistic clarification regarding its functional relevance (Hiyoshi *et al*, 2009; Ma *et al*, 2019).

Notably, several inflammatory CKs act as potent pro-fibrotic mediators: examples with implications in liver are CCL2, CCL20 and CCL24 (Chu *et al*., 2018; Govaere *et al*., 2020; Greenman *et al*., 2023; Hanson *et al*., 2019; Xi *et al*., 2021). The CXCL12/CXCR4 axis promotes leukocyte recruitment and hepatic stellate-cell activation in injured liver, yet, pharmacological targeting of CXCR4 has produced both protective and detrimental outcomes *in vivo*, underscoring the context-dependent nature of CXCR4 signaling in liver fibrosis (Boujedidi *et al*, 2015; Hong *et al*, 2009; Saiman *et al*, 2015; Sung *et al*, 2018; Wald *et al*, 2004).

Here, we set out to identify chemokine interactions of MIF-2. An unbiased chemokine array with SPR validation revealed that MIF-2 displays a broader chemokine interaction spectrum than MIF (Brandhofer *et al*., 2022; Eckardt *et al*., 2020; von Hundelshausen *et al*., 2017). Whereas reported MIF interactions include CXCL4L1, CXCL9, CCL28, Prdx1, and Prdx6, the MIF-2 interactome encompassed CCL20, CCL24–26, CCL28, CXCL9, CXCl11, CXCL13, CXCL17, CCL21, CXCL12α, XCL1, and Prdx1. Thus, MIF and MIF-2 share only a limited set of interactors, namely CCL28, CXCL9, and Prx1, and CCL20 emerging as a specific interactor of MIF-2.

Given the prominent hepatic expression and metabolic activity of MIF-2, we next examined which newly identified MIF-2 interactors might be most relevant in hepatic fibrosis (El Bounkari *et al*., 2025; Gilgenkrantz *et al*., 2025). CCL20 emerged as a strong candidate based on its established role in liver fibrosis and inclusion in the fibrosis-progression signature according to Govaere et al (Chu *et al*., 2018; Govaere *et al*., 2020; Hanson *et al*., 2019). To contextualize MIF-2 and its receptor network during fibrotic progression, we re-analyzed their data set, comprising 206 human liver biopsies spanning the full histological spectrum from NAFL and NASH with fibrosis stages F0–F4. CCL20 showed stage-dependent induction along disease progression, whereas MIF and MIF-2 were constitutively expressed across all stages (Govaere *et al*., 2020; Rinella *et al*, 2023). CD74 and CXCR4 were abundantly expressed in the liver, but only CXCR4 increased in a stage-dependent manner, consistent with enhanced influx of CXCR4⁺ leukocytes during fibrotic progression (Boujedidi *et al*., 2015; Saiman *et al*., 2015; Sung *et al*., 2018; Wald *et al*., 2004). In contrast, CCR6 — the only known receptor for CCL20 — remained low and unchanged across disease stages. This divergence between strong CCL20 induction and minimal, stable CCR6 expression suggests limited canonical CCL20 signaling in fibrotic liver and raises the possibility that CCL20 exerts alternative functions, including modulation of CXCR4 signaling via MIF-2/CCL20 complex formation. In line with this concept, a CCR6-independent CCL20–integrin α5β1 interaction was shown to promote fibroblast activation in pulmonary fibrosis (Liu *et al*., 2025). Among the remaining array-positive CKs, only CCL21 and CCL28 increased with fibrosis, identifying them as candidates for future studies.

Beyond chemokine array and SPR analyses, MIF-2/CCL20 complex formation was validated by MST, peptide array mapping, and AlphaFold 3.0 modeling. Peptide array identified two MIF-2–responsive regions within CCL20, comprising an N-terminal β-sheet–associated stretch adjacent to the CC-motif and a C-terminal binding region. Consistent with this, the cryo-EM structure of the CCL20–CCR6 complex indicates that the CCL20 β-sheet surface engages CCR6 extracellular loops, while the N-terminus inserts into the orthosteric pocket and β-sheet-CK-CK interactions are known to modulate receptor activation (Koenen *et al*, 2009; von Hundelshausen *et al*., 2017; Wasilko *et al*., 2020). AlphaFold modeling further suggested docking CCL20 C terminus into the MIF-2 tautomerase pocket, in line with the observed inhibition of MIF-2 tautomerase activity by CCL20 to a degree comparable to 4-CPPC (Pantouris *et al*., 2018; Tilstam *et al*., 2019).

The co-occurrence of high basal MIF-2 expression and fibrosis-associated CCL20 induction prompted analysis of protein expression in human liver tissue. MIF-2 and CCL20 protein levels correlated across samples, suggesting coordinated regulation, although CCL20 protein levels did not differ between non-fibrotic and fibrotic tissue despite transcriptomic induction. Pulldown assays and PLA confirmed MIF-2/CCL20 complex formation in human liver tissue, with increased complex detection in fibrotic compared with non-fibrotic samples (Alam, 2022). These findings suggest fibrosis-associated redistribution or compartmental enrichment of the two proteins, potentially through increased extracellular release, matrix anchoring, or localization within inflammatory niches. However, given the limited number of human samples, these observations should be considered hypothesis-generating and require validation in larger cohorts, including analyses stratified by relevant clinical variables.

We next examined the functional consequences of MIF-2/CCL20 complex formation in settings relevant to hepatic inflammation and fibrosis. CXCR4-driven T-cell trafficking to the liver is well established, and our transcriptomic analyses revealed increased CXCR4 expression during fibrosis, while the CCL20–CCR6 axis also contributes to hepatic T-cell recruitment (Boujedidi *et al*., 2015; Wang *et al*., 2024b). In a tissue-mimicking 3D chemotaxis assay, MIF-2 induced robust CD4⁺ T-cell migration, identifying MIF-2 as a previously unrecognized T-cell chemoattractant, whereas CCL20 alone elicited only a weak, non-significant response, likely reflecting low CCR6⁺ expression of CD4⁺ T cells. Notably, combined MIF-2/CCL20 stimulation completely abolished chemotaxis, suggesting that complex formation limits excessive T-cell accumulation, fine-tunes the balance of relevant T-cell phenotypes, or acts as a stop signal that terminates T-cell movement within fibrotic niches. To determine whether this also affects stromal effector functions, we examined IL-6 secretion in fibroblasts as a second fibrosis-related readout. Consistently, while MIF-2 or CCL20 alone induced fibroblast IL-6 secretion, combined stimulation markedly suppressed IL-6 production. IL-6 promotes α-SMA expression and myofibroblast differentiation in fibroblasts, and reduced IL-6 secretion under MIF-2/CCL20 co-stimulation may therefore limit progression toward a profibrotic phenotype, although myofibroblast differentiation was not assessed in this study (Gallucci *et al*., 2006). In line with this concept, MIF-2 has previously been reported to enhance fibroblast proliferation and viability (Kim *et al*., 2017). Elevated IL-6 levels correlate with liver disease severity, fibrosis and mortality, yet IL-6 exerts both regenerative and profibrotic functions depending on disease context and signaling mode (Cheng *et al*, 2022; Hou *et al*, 2021; Kar *et al*, 2019; Remmler *et al*, 2018; Xiang *et al*, 2018). In acute liver injury, IL-6 promotes hepatocyte survival and regeneration via gp130/STAT3, and IL-6-deficient mice show impaired repair responses (Blindenbacher *et al*, 2003; Cheng *et al*., 2022; Cressman *et al*, 1996). In chronic disease contexts, IL-6 has been implicated in hepatic stellate-cell (HSC) activation, and upregulation of IL-6–STAT3 signaling has been observed in fibrotic lesions (Kagan *et al*, 2017; Xiang *et al*., 2018). The inhibitory effect of MIF-2/CCL20 on fibroblast IL-6 production suggests that complex formation modulates stromal cytokine release and the local cytokine milieu within fibrotic niches. Such chemokine–cytokine crosstalk may contribute to the context-dependent activities of inflammatory mediators in chronic liver disease. A limitation of this study is the use of dermal fibroblasts rather than liver-resident mesenchymal cells. In the fibrotic liver, activated HSCs represent the principal source of myofibroblasts, while portal fibroblasts form a smaller but functionally relevant proliferative pool (Lei *et al*, 2022; Mederacke *et al*, 2013). Accordingly, future studies should validate these findings in HSCs or other liver-resident cell populations.

This study has further limitations. Although our data indicate that MIF-2/CCL20 complex formation can influence CXCR4-dependent responses, its contribution to immune-cell recruitment *in vivo* remains unresolved, and dedicated liver injury models will be required to clarify whether this interaction exerts pro- or anti-fibrotic effects. Moreover, the precise molecular mechanism underlying the observed inhibition remains incompletely defined and may involve additional regulatory modes, including negative cooperativity, receptor heterodimerization, or functional crosstalk (Stephens & Handel, 2013). While the MIF/CD74 axis has been reported to exert anti-fibrotic effects in liver fibrosis, we did not address whether MIF-2/CCL20 complex formation modulates CD74-dependent signaling (Heinrichs *et al*., 2011).

Together, our study advances the emerging concept of ACK–CK interactions by characterizing the MIF-2 chemokine interactome and identifying MIF-2/CCL20 complex formation as an intrinsic modulator of inflammatory and fibrotic signaling. We provide a mechanistic framework to explain context-dependent chemokine activities in chronic liver disease. Addressing a current gap in fibrosis- and chemokine-directed therapeutic strategies, our findings suggest that a deeper understanding of endogenous MIF-2/CCL20 complex formation could prove pivotal for the development of MIF-2– or CCL20-targeting inhibition approaches in inflammation and fibrosis.

## Methods

### Proteins and reagents

Recombinant human MIF-2 was produced as previously described with minor modifications (El Bounkari *et al*., 2025; Merk *et al*., 2011). The MIF-2 coding sequence was cloned into a pET22b vector, expressed in E. coli BL21-CodonPlus (DE3) cells, and purified from clarified lysates by anion-exchange chromatography on a Q Sepharose column (Cytiva, Freiburg, Germany) using fast protein liquid chromatography (FPLC; Cytiva), followed by high-performance liquid chromatography on a C18 reverse-phase column (RP-HPLC; Cytiva) to achieve final purification. Refolding by stepwise dialysis yielded recombinant MIF-2 of approximately 98% purity and <10 pg LPS/µg protein, as determined using a recombinant Factor C assay (Cambrex, East Rutherford, USA).

MST-Red–MIF-2 for MST experiments was generated by labeling recombinant MIF-2 using a Monolith Protein Labeling Kit RED-NHS, second generation (NanoTemper, Munich, Germany), according to the manufacturer’s instructions. Biotinylated MIF-2 was produced using a Biotin Protein Labeling Kit (Roche, Mannheim, Germany) with either Biotin-7-NHS or biotin-amidohexanoic acid N-hydroxysuccinimide ester (Sigma-Aldrich, St. Louis, MO, USA). Recombinant human PRDX1 and PRDX6 (Abcam PLC, Cambridge, UK), HBD-1 and HBD-2 (ProSpec-Tany TechnoGene Ltd., Ness Ziona, Israel), HMGB1 (Novus Biologicals Europe, Abingdon, UK), CXCL4L1 and CXCL4 (ChromaTec, Greifswald, Germany), and all other recombinant human chemokines (Peprotech, Hamburg, Germany) were used as indicated.

### Chemokine protein array

The chemokine protein array was performed as previously described for MIF, using biotinylated human MIF-2 instead (Brandhofer *et al*., 2022). Recombinant human CKs and selected ACKs were spotted on nitrocellulose membranes at 100 ng per spot. Membranes were blocked with 1× ROTI®Block (Carl Roth, Karlsruhe, Germany) for 2 h at room temperature and incubated overnight with biotin-MIF-2 (1 µg/mL) in 10 mM Tris-HCl (pH 8.0) or 10 mM MES (pH 6.0). After three washes with 0.01% Tween 20 in water, membranes were incubated for 2 h with horseradish-peroxidase–conjugated streptavidin (Bio-Techne GmbH, Wiesbaden, Germany; 1:200 in 1× ROTI®Block). Bound biotin-MIF-2 was detected by chemiluminescence using SuperSignal™ West Pico substrate (Thermo Fisher, Waltham, MA, USA) and imaged with a LAS-3000 Imaging System (Fuji Photo Film Co., Ltd., Japan).

### Surface plasmon resonance (SPR) analysis

The surface plasmon resonance experiments were performed using a Biacore X100 (Cytiva, Freiburg, Germany). Biotinylated MIF-2 was immobilized on a neutravidin-coated C1 chip at a density of approximately 1065 Response Units on flow cell 2. Recombinant human chemokines (CCL2, CCL20, CCL24, CCL25, CCL26, CCL28, CXCL9, CXCL11, CXCL12, CXCL13, and CXCL17; all from PeproTech) in running buffer (HBS-EP⁺) were injected at various concentrations as indicated in Figure 2 at a flow rate of 60µL/min. Association and dissociation phases were recorded for 60-180 s and 240 s, respectively. In two instances (i.e., CCL2 and CXCL9), single cycle kinetics was performed as total regeneration was difficult to achieve. Sensor surfaces were regenerated with two 60-s pulses of 30 mM NaOH and 2 M NaCl. Binding data were fitted using a 1:1 Langmuir interaction model in the Biacore X100 Evaluation Software 2.1.0.201 Plus (Cytiva).

### Re-analysis of publicly available RNA-sequencing data

Gene-level expression values normalized as transcripts per million (TPM) and corresponding sample metadata were obtained from the publicly available discovery cohort of Govaere et al. (GEO accession GSE135251), comprising 206 human liver biopsies spanning the full histological spectrum of NAFLD from NAFL (n = 51) to NASH with fibrosis stages F0–F4 (n = 34, 53, 54, and 14, respectively) (Govaere *et al*., 2020). Gene expression analyses were performed in Python (v3.11.11) using pandas v.2.2.3, numpy v.2.1.3, and matplotlib v.3.10.1. Gene identifiers were mapped to HGNC symbols using the GRCh38.p13 annotation file. Expression patterns of MIF, MIF-2 (DDT), CCL20, their receptors (CXCR4, CCR6, CD74), and additional chemokines and mediators (CCL21, CCL24, CCL25, CCL26, CCL28, CXCL9, CXCL11, CXCL12, CXCL13, CXCL17, XCL1, and PRDX1) were evaluated across disease stages. Expression values were log1p transformed and visualized using boxplots with overlaid strip plots generated with seaborn v.0.13.2. Statistical comparisons were performed using the Mann–Whitney U test via the stat-annotations package v.0.7.1 with Holm–Bonferroni correction. Analysis scripts are available at https://github.com/SimonE1220/MIF2_CCL20_BulkSeq_Liver.

### Microscale thermophoresis (MST)

MIF-2–CCL20 protein interactions were analyzed by MST using a Monolith NT.115 instrument equipped with green/red filters (NanoTemper). Measurements were performed at 25 °C with MST power settings of 40% and 80% and LED excitation adjusted to 90–95% to yield initial fluorescence counts of approximately 700–800. Thermophoresis was recorded for 40 s (−5 to +35 s), with sample heating applied from 0 to 30 s. MST-Red–labeled MIF-2 was mixed 1:1 with serial dilutions of CCL20 (PeproTech) in assay buffer (10 mM Tris–HCl, pH 8.0, 0.01% BSA), resulting in final MIF-2 concentrations of 100 nM, and incubated on ice for at least 30 min prior to measurement. Data from independent experiments were analyzed using MO.AffinityAnalysis V2.3 software (NanoTemper) with default KD model and T-jump settings by evaluating the temperature-related intensity change between cold (−1 to 0 s) and hot (0.5 to 1.5 s) regions. Curve fitting was performed in GraphPad Prism v9 using a one-site total binding model.

### CelluSpot peptide array

An established cellulose-based peptide array approach was used to map MIF-2 binding sites within the CCL20 sequence (Brandhofer *et al*., 2022; Lacy *et al*, 2018). Overlapping 15-mer peptides spanning the complete human CCL20 sequence were synthesized with a three–AA offset, resulting in a 12–residue overlap between adjacent peptides, using the MultiPep RSi/CelluSpot Array system (Intavis, Cologne, Germany). Peptides were produced as cellulose membrane–bound spots, released by membrane dissolution, and transferred onto coated glass slides using an automated Intavis microarrayer. Arrays were equilibrated in blocking buffer (50 mM Tris-buffered saline, pH 7.4, 0.1% Tween 20, supplemented with 1% BSA), washed in the same buffer without BSA, and incubated with biotinylated human MIF-2 (3 µM in blocking buffer). Bound MIF-2 was detected using streptavidin–horseradish peroxidase (Roche, Mannheim, Germany) diluted in blocking buffer, followed by chemiluminescent detection with SuperSignal West Dura substrate and imaging on an Odyssey Fc system.

### *In-silico* protein–protein interaction modeling

Protein–protein interaction models were generated using AlphaFold 3.0 with human MIF-2/DDT (UniProt ID: P30046) and mature CCL20 (UniProt ID: P78556) as input sequences. The initiator methionine of MIF-2 and the N-terminal signal peptide of CCL20 were removed to model the biologically active, secreted forms. Predicted heterodimeric models and interface regions were visualized and analyzed in PyMOL (v3.2.0a)

### MIF-2 tautomerase activity assay

MIF-2 tautomerase activity was quantified using the classical 4-hydroxyphenylpyruvate (HPP) tautomerization assay. Reactions (1 mL) were performed at RT in 0.5 M boric acid buffer (pH 6.2; Sigma-Aldrich, Cat. No. B0394-500G) containing recombinant MIF-2 (55 nM), either the MIF-2 inhibitor 4-CPPC (30 nM) or recombinant human CCL20 (30 nM; PeproTech), and HPP (BCCJ, Cat. No. 114286-5G). For competition experiments, MIF-2 was pre-incubated with 4-CPPC or CCL20 for 20 min prior to substrate addition. Tautomer formation was monitored by measuring absorbance at 306 nm for 300 s. Tautomerase activity was calculated from the initial linear phase of the reaction using an experimentally determined molar extinction coefficient (ε₃₀₆) obtained under the same assay conditions. Initial velocities and AUCs (0–120 s) were calculated from baseline-corrected kinetic traces, normalized to MIF-2 alone, and reported as mean ± SD from triplicate measurements. Data analysis was performed using Origin 2024.

### Human liver tissue specimens

Double-coded human liver tissue specimens and corresponding clinical data used in this study were provided by the Biobank of the Department of General, Visceral and Transplant Surgery at Ludwig-Maximilians-University (LMU) Munich, operating under the administration of the Human Tissue and Cell Research (HTCR) Foundation, whose framework—including the acquisition of written informed consent from all donors—has been approved by the ethics commission of the Faculty of Medicine at LMU Munich (ethics vote 025-12) and by the Bavarian State Medical Association (approval number 11142), with the present study additionally approved by the LMU Munich ethics committee (ethics vote 21-1187). Clinically annotated patient characteristics, including fibrosis grading assessed on matched histological sections by a board-certified pathologist, are summarized in Supplementary Table 1. Snap-frozen liver tissue and cryoconserved sections were used for the protein-based and imaging-based assays described below.

### SDS-PAGE and Western blot

For Western blot analysis, frozen human liver samples were lysed in Pierce™ RIPA Lysis and Extraction Buffer (Thermo Fisher) and homogenized using a TissueLyser LT (Qiagen, Hilden, Germany). Lysates were clarified by centrifugation, and protein concentrations were determined using the Pierce™ BCA Protein Assay Kit and an EnSpire Multimode Plate Reader (PerkinElmer, Waltham, MA, USA). Samples were diluted in LDS sample buffer (NuPAGE™, Thermo Fisher), heated at 95 °C for 15 min, and equal protein amounts were separated on 15% SDS–PAGE gels (NuPAGE™, Thermo Fisher) and transferred to nitrocellulose membranes. The CozyHi Prestained Protein Ladder (highQu GmbH, Kraichtal, Germany) served as the molecular weight marker. Membranes were blocked for 1 h at RT in PBS–Tween containing 5% nonfat dry milk and incubated overnight at 4 °C with primary antibodies against β-actin (C4, sc-47778, 1:1000; Santa Cruz, Dallas, TX, USA), MIF-2 (rabbit polyclonal antibody (Merk *et al*., 2011), 1:1000), or CCL20/MIP-3α (LS-B7409, 1:250; LSBio, Seattle, WA, USA). After washing, membranes were incubated with HRP-conjugated secondary antibodies (1:10 000; Thermo Fisher), and signals were detected using SuperSignal™ West Dura substrate and recorded on an Odyssey® Fc Imager (LI-COR Biosciences GmbH, Bad Homburg, Germany).

### MIF-2–CCL20 pulldown assays

For semi-endogenous pull-down experiments, human liver lysates were prepared as described for WB analysis, diluted 1:1 with ice-cold PBS, and aliquots were retained as input controls. All incubation steps were performed on a rotating wheel at 4 °C unless otherwise indicated. Lysates were incubated for 2 h with or without recombinant His-tagged human CCL20 (His–CCL20; Sino Biological, Beijing, China), followed by ON capture using pre-washed His-tag–binding magnetic beads (Dynabeads™ His-Tag Isolation and Pulldown, Thermo Fisher) to capture His–CCL20 together with associated endogenous binding partners. Beads were collected and washed three times with PBS containing 0.1% Tween-20.

To complement this semi-endogenous approach, a reciprocal pull-down was performed using biotinylated human MIF-2 immobilized on streptavidin-coated magnetic beads (Dynabeads™ M-280 Streptavidin, Thermo Fisher) to assess direct interaction with recombinant CCL20. After immobilization in binding buffer (10 mM Tris-HCl, pH 8.0, 0.1% BSA), beads were incubated ON with recombinant human CCL20 (PeproTech, Rocky Hill, NJ, USA), with buffer-only beads serving as controls. Eluted proteins from both approaches were denatured in LDS sample buffer at 95 °C and subjected to SDS–PAGE/WB as described above.

### Immunofluorescent co-staining and proximity ligation assay (PLA) in human liver tissue

Immunofluorescent (IF) co-stainings and *in situ* proximity ligation assays (PLA) were performed on human liver cryosections obtained from the LMU HTCR biobank. Sections were briefly warmed to 37 °C, air-dried for 30 min at RT, fixed and permeabilized in cold acetone (Merck, Darmstadt, Germany) for 6 min at 4 °C, air-dried for 30 min, and rehydrated in PBS (Thermo Fisher). All samples were processed identically up to this step before IF or PLA.

For IF staining, sections were blocked for 1 h at RT in 1% BSA in PBS (Carl Roth) and incubated ON at 4 °C with primary antibodies against D-DT/MIF-2 (rabbit polyclonal, 1:300, (Merk *et al*., 2011)) and CCL20 (goat anti-CCL20, PA5-47517, Thermo Fisher). After washing, fluorophore-conjugated secondary antibodies were applied for 1 h at RT (donkey anti-rabbit Cy3, 1:300, Jackson ImmunoResearch, West Grove, PA, USA; donkey anti-goat Alexa Fluor 647, 1:500, Thermo Fisher), followed by DAPI counterstaining (Sigma-Aldrich) and mounting with Vectashield Antifade Mounting Medium (Vector Laboratories, Newark, CA, USA).

PLA for detection of MIF-2/CCL20 complexes was performed using the Duolink™ In Situ Orange Starter Kit Goat/Rabbit (DUO92106-1KT, Sigma-Aldrich) according to the manufacturer’s instructions, using the same primary antibodies as for IF. After blocking for 1 h at 37 °C, sections were incubated ON at 4 °C with primary antibodies, followed by incubation with PLUS and MINUS PLA probes (1:5) for 1 h at 37 °C, ligation for 30 min at 37 °C, and signal amplification for 100 min at 37 °C. Sections were washed using Duolink® Wash Buffers A and B and mounted with Duolink® Mounting Medium with DAPI (82040-0005, Sigma-Aldrich). Across samples, minor tissue autofluorescence was detected predominantly in the Cy5 channel with partial spillover into the Cy3 channel. Because true PLA signal was confined to the Cy3 channel, Cy5 autofluorescence was quantified and subtracted to prevent false-positive signal detection.

Imaging for both IF and PLA was performed using a Leica DMi8 fluorescence microscope (Leica, Wetzlar, Germany) or a Zeiss LSM880 AiryScan confocal microscope (Carl Zeiss, Jena, Germany), using identical exposure or laser settings across samples. PLA signals were quantified by counting PLA signals in ≥4 randomly selected fields per specimen using automated analysis in Fiji (ImageJ, version 2.16.0).

### Isolation of primary human CD4⁺ T cells from peripheral blood mononuclear cells

Peripheral blood mononuclear cells (PBMCs) were isolated from leukocyte-enriched peripheral blood obtained from thrombocyte apheresis filters (Leukoreduction System, LRS chambers) of anonymous healthy donors at the Department of Transfusion Medicine, LMU University Hospital by density-gradient centrifugation using Ficoll-Paque Plus (GE Healthcare). Residual erythrocytes were removed by incubation in RBC Lysis Buffer (BioLegend, San Diego, CA, USA), followed by washing in RPMI 1640 supplemented with 10% fetal bovine serum (FBS). CD4⁺ T cells were purified from PBMCs by magnetic negative selection using the Human CD4⁺ T Cell Isolation Kit (Miltenyi Biotec, Bergisch Gladbach, Germany). Purity routinely exceeded 95%, as assessed by flow cytometry using anti-CD3 and anti-CD4 antibodies (Zhang *et al*., 2024). Purified CD4⁺ T cells were rested ON in RPMI 1640 supplemented with 10% FBS at 37 °C and 5% CO₂ prior to use. All procedures complied with the Declaration of Helsinki and were approved by the Ethics Committee of LMU Munich (approvals 18-104 and 23-0639)

### 3D chemotaxis assay

3D migration of human CD4⁺ T cells was analyzed using the µ-Slide Chemotaxis system (Ibidi GmbH, Gräfelfing, Germany) as previously described (Brandhofer *et al*., 2022; Hoffmann *et al*, 2020). Rested CD4⁺ T cells were embedded at 4 × 10⁶ cells per gel in rat tail collagen type I (Ibidi) polymerized in DMEM within the central observation channel. Stable or competitive gradients were generated by loading the reservoirs with control medium, recombinant human MIF-2 (400 ng/ml), CCL20 (300 ng/ml), or both chemokines prepared in RPMI 1640.

Time-lapse imaging was performed at 37°C for 90 min on a Leica DMi8 inverted microscope equipped with a DMC2900 CMOS digital camera, with images acquired at 1-min intervals. Cell trajectories were reconstructed in ImageJ using the Manual Tracking plugin and analyzed with the Ibidi Chemotaxis and Migration Tool.

### Fibroblast IL-6 secretion assay

Primary human foreskin dermal fibroblasts (HFDFs, passages 6–9) were cultured in DMEM supplemented with 10% FBS and 1% penicillin–streptomycin at 37 °C and 5% CO₂. For stimulation, cells were washed and incubated in serum-free DMEM and treated for 16 h with recombinant human MIF-2 (1000 ng/mL), recombinant human CCL20 (500 ng/mL; PeproTech), or both combined, with chemokines pre-incubated together for 30 min at RT prior to addition. Supernatants were collected and clarified by centrifugation.

Human IL-6 concentrations were quantified using the Human IL-6 DuoSet ELISA (DY206; R&D Systems, Minneapolis, MN, USA) according to the manufacturer’s instructions and measured at 450 nm with wavelength correction at 540–570 nm using an EnSpire Multimode Plate Reader (PerkinElmer). Concentrations were calculated from plate-specific standard curves.

### Flow cytometry

Cell-surface expression of CXCR4, CD74, and CCR6 on HFDFs and CCR6 on isolated human CD4⁺ T cells was analyzed by flow cytometry using fluorophore-conjugated antibodies and appropriate isotype controls. Antibodies included APC-conjugated anti-CXCR4 (BioLegend; Cat# 306510), FITC-conjugated anti-CD74 (BD Biosciences; Cat# 555540), and anti-CCR6 antibodies (PE-conjugated, Miltenyi; Cat# 130-100-377; APC/Cy7-conjugated, BioLegend; Cat# 353432). Cells (2 × 10⁵) were stained for 1 h at 4 °C in the dark in PBS containing 0.5% BSA and analyzed on a BD FACSVerse or BD FACSSymphony A1. Data were processed using FlowJo v10.2 (Tree Star, Ashland, OR, USA).

### Statistical analysis

Statistical analyses were performed using GraphPad Prism version 9 (GraphPad Software, San Diego, CA, USA). Data are presented as means ± standard deviation (SD). Normality was assessed using the Shapiro–Wilk test together with inspection of QQ plots to confirm distributional assumptions. Group comparisons were performed using unpaired two-tailed Student’s t-tests or one-way ANOVA with Tukey’s post hoc correction. A *P* value < 0.05 was considered statistically significant

### Data availability and material

All data and materials as well as software application information are available in the manuscript, the supplementary information, or are available from the corresponding authors upon reasonable request. The bulk-RNAseq dataset published by Govaere et al., which was re-analyzed during the current study is publicly available on the gene expression omnibus (GEO) under accession number GSE135251 (24). The full analysis code is published on GitHub: https://github.com/SimonE1220/MIF2_CCL20_BulkSeq_Liver.

## Authors’ contributions

A.H., M.B., P.S. and J.B. conceived and designed the study. A.H., M.B., G.B., Y.H., E.S., M.K.O., X.B., L.Z., M.H. S.E., C.K., K.H. performed research and analyzed data. A.H., M.B., O.E.B., A.K., P.v.H., P.S. and J.B. contributed to the interpretation of the data. C.W., H.N. and A.K. contributed critical materials and methods. The first draft of the manuscript was written by A.H. with help from P.S. and J.B.; all authors critically reviewed, revised and commented on the manuscript drafts and approved the final draft. A.H., P.v.H., A.K. and J.B. provided funding for the study.

## Acknowledgements

This work was supported by Deutsche Forschungsgemeinschaft (DFG) grant SFB1123-A3 to J.B. and A.K., as well as German Center for Cardiovascular Diseases (DZHK) grant FKZ 81X2600278, DZHK B 23-008 Ext to J.B. A.H. was supported by a Metiphys scholarship of LMU Munich, funding by the Knowledge Transfer Fund (KTF) of the LMU Munich-DFG excellence (LMUexc) program, by the Friedrich-Baur-Foundation e.V. and the Association to Promote Science and Research (WiFoMed) of the LMU Faculty of Medicine. P.v.H. and J.B. acknowledge support from DFG / SFB1123-A2. C.Wi is supported by a research grant of the Else Kröner-Fresenius-Stiftung (2021_EKFS.92). C.We. acknowledges support from DFG projects SFB1123-A1 and A10. This study was supported by the Human Tissue and Cell Research Foundation, a nonprofit foundation regulated by German civil law, which facilitates research with human tissue through the provision of an ethical and legal framework for (prospective/retrospective) sample collection. We thank Maida Avdic/Simona Gerra for excellent technical support and Bernhard Zwissler for the provision of further personnel funding to A.H.; ChatGPT (OpenAI) was used to assist in shortening original text for this manuscript. All final revisions and content have been reviewed and approved by the authors.

